# Beta Desynchronization Mediates The Impact Of Internally Directed Attention In A Dual-Task Working Memory Paradigm

**DOI:** 10.1101/2025.07.23.666304

**Authors:** Ankit Yadav, Arpan Banerjee, Dipanjan Roy

## Abstract

Internally directed attention (IDA), whether spontaneous or intentional, has been associated with impaired performance on externally directed cognitive tasks. Yet, the neurophysiological mechanisms underpinning this disruption remain poorly understood. In the present study, we characterized the neural correlates of IDA and identified how they impact on performance in a color-recall working memory task using electroencephalography (EEG). Participants performed a novel dual-task paradigm involving either self-referential (IDA) or perceptual processing of adjectives, involving externally directed attention (EDA) followed by a color-recall task. IDA enhanced late positive potentials (LPP) in EEG over medial-frontal electrodes as a marker of sustained affective and self-referential engagement. Time–frequency analyses further revealed increased event-related desynchronization in alpha and beta bands during stimulus encoding in the IDA condition, as well as increased alpha synchronization during the delay period, consistent with internal attention maintenance. To capture trial-level variability in task performance, we applied conditional quantile regression on the single trial data. Results showed that beta desynchronization in interaction with condition type during encoding influenced performance significantly in trials with low errors, whereas trials with high error in color recall were better explained by increased reaction times. These findings provide converging electrophysiological evidence for distinct neural signatures of internally directed attention and highlight their behavioral consequences in working memory performance.

## Introduction

Attention continuously shifts between the external world, for example, counting something on the screen, and the internal world, reflecting on feelings about own mental state shaped by last night’s dinner. Complex cognitive functions, such as working memory, rely on such seamless interplay between externally directed attention (EDA) and internally generated thought processes. According to Dixon et al. (2014), internally directed cognition in turn involves internally directed attention (IDA) directed towards information in long-term memory, working memory, prospective and retrospective thinking, and self-referential content. This includes episodic memory retrieval, simulating future events, stimulus-independent thoughts, mental imagery, and dreaming. IDA is largely decoupled from the external environment, operating independently or in response to internal or external stimuli (see Chun et al., 2011 for review). In contrast, EDA refers to the attention directed towards the external perceptual world and involves processing sensory information in a specific modality, such as visual and auditory input (Chun et al., 2011; Dixon et al., 2014).

While internally focused states are essential for introspection and self-evaluation, when we have an EDA task at hand, switching to such states during encoding and maintenance phases of the task may detract the cognitive resources necessary for processing and encoding externally stimuli, ultimately impairing performance on the task that require sustained attention (Kam & Handy, 2014; Kane & McVay, 2012; Killingsworth & Gilbert, 2010; Rummel & Boywitt, 2014; Smallwood et al., 2008). Self-referential processing (SRP), a central aspect of IDA, plays a crucial dual role in cognition by shaping subjective experience and modulating task performance by promoting internally generated thoughts—often at the expense of task-focused attention (Dixon et al., 2014; Huijser et al., 2018).

Neuroimaging studies show SRP not only recruits the default mode network (DMN) and cortical midline structures but also interacts dynamically with attention and control networks (Benedek et al., 2016; Dixon et al., 2014). The balance between EDA and IDA is mediated by the interplay among the DMN, which supports internally generated thought, and frontoparietal control and dorsal attention networks, which support goal-directed and perceptual processing. When self-relevance is high, SRP can shift this balance toward internally focused cognition (Raichle et al., 2001; Zhao et al., 2018).

Beyond neuroimaging evidence, event-related potential (ERP) studies have consistently reported enhanced late positive potentials (LPP) in response to self-relevant (Katyal et al., 2020) and emotionally arousing stimuli (De Cesarei & Codispoti, 2011; Hajcak et al., 2009; Naumann et al., 1992; Schindler et al., 2023; Schupp et al., 2000). This pattern suggests that, compared to EDA, conditions promoting IDA are associated with sustained affective engagement with the stimulus content. In oscillatory dynamics, particularly in alpha and theta band, differential processing has been reported in IDA and EDA (Kam et al., 2018; Kam & Handy, 2014; Mu & Han, 2010; Subramaniam et al., 2019). IDA is associated with increased power in the alpha frequency band (Cooper et al., 2003, 2006; Kam et al., 2018; Mu & Han, 2010; O’Connell et al., 2009; Schupp et al., 1994), while EDA is accompanied by an increase in power in the theta frequency band (Kam et al., 2018, 2022; Hammer et al., 2024). Research on attentional reorientation suggests that beta power increase plays a role in redirecting attention from internal distractions (e.g., stress-induced intrusive thoughts) to external tasks (Palacios-García et al., 2021), while beta desynchronization is associated retrieval of self-generation information (Graber & Fujioka, 2020; Subramaniam et al., 2019).

Despite these advances, some aspects of IDA on complex cognitive tasks remain unexplored. Specifically, it is unclear whether IDA influences the encoding of the perceptual features of the stimuli when cognitive resources are limited, or whether it impacts the recall by reducing rehearsal during the maintenance period. Moreover, although prior research has identified EEG signatures associated with IDA, it is still not known which of these neural features specifically drive its detrimental impact on task performance. Considering this, the current study employed a novel color-recall working memory paradigm that juxtaposed self-referential processing guided by IDA with an analytical condition involving EDA while EEG data were recorded. First, a paradigm was established that effectively induced differential processing in IDA and EDA conditions. Encoding the stimuli in self-reference would lead to IDA was the key hypothesis of the study. This, in turn, was expected to lead to enhanced LPP amplitude relative to EDA and relatively reduced power in alpha and beta band activity. While in the delay period, IDA was hypothesized to result in increased activity in the alpha band over posterior sensors. The focus was on analyzing the behavioral and electrophysiological features that influenced different levels of task performance, using quantile regression to examine the trial-to-trial variability that had often been missed by earlier studies. By linking IDA’s oscillatory signatures to working memory performance, this study aimed to advance our understanding of how internal thought shapes goal-directed behavior.

## Materials and Methods

### 2.1 Participants

Thirty participants (16 females and 14 males, mean age = 25.2 years, SD = 2.3 years) participated in a study involving reporting perceptual behavior and simultaneous electroencephalogram (EEG) recordings. Participants were required to have a university degree with English as the medium of instruction. All the participants reported normal or corrected-to-normal vision, no history of color-blindness, were right-handed, and declared no history of neurological or psychiatric disorders. Participants were compensated monetarily for their time nominally and gave informed consent to participate in format approved by Institutional Human Ethics Committee (IHEC) of National Brain Research Centre (NBRC), India. EEG data from two participants were excluded because the electrode impedance exceeded the set threshold when checked at the end of the recording session.

### 2.2 Behavioral paradigm

A complex color-recall working memory task was implemented (**Figure 1** illustrates the trial sequence). At the start of each trial, a cue word appeared in the centre of the screen for a variable duration, randomly sampled from a discrete uniform duration between 0.7 s and 1.2 s ms in 0.1 s increment (mean = 0.97 s, SD = 0.17 s). The cue indicated which task the participant should perform:

- In the EDA condition, the cue word was *“vowel”*.
- In the IDA condition, the cue word was *“self”*.

**Figure 1.**
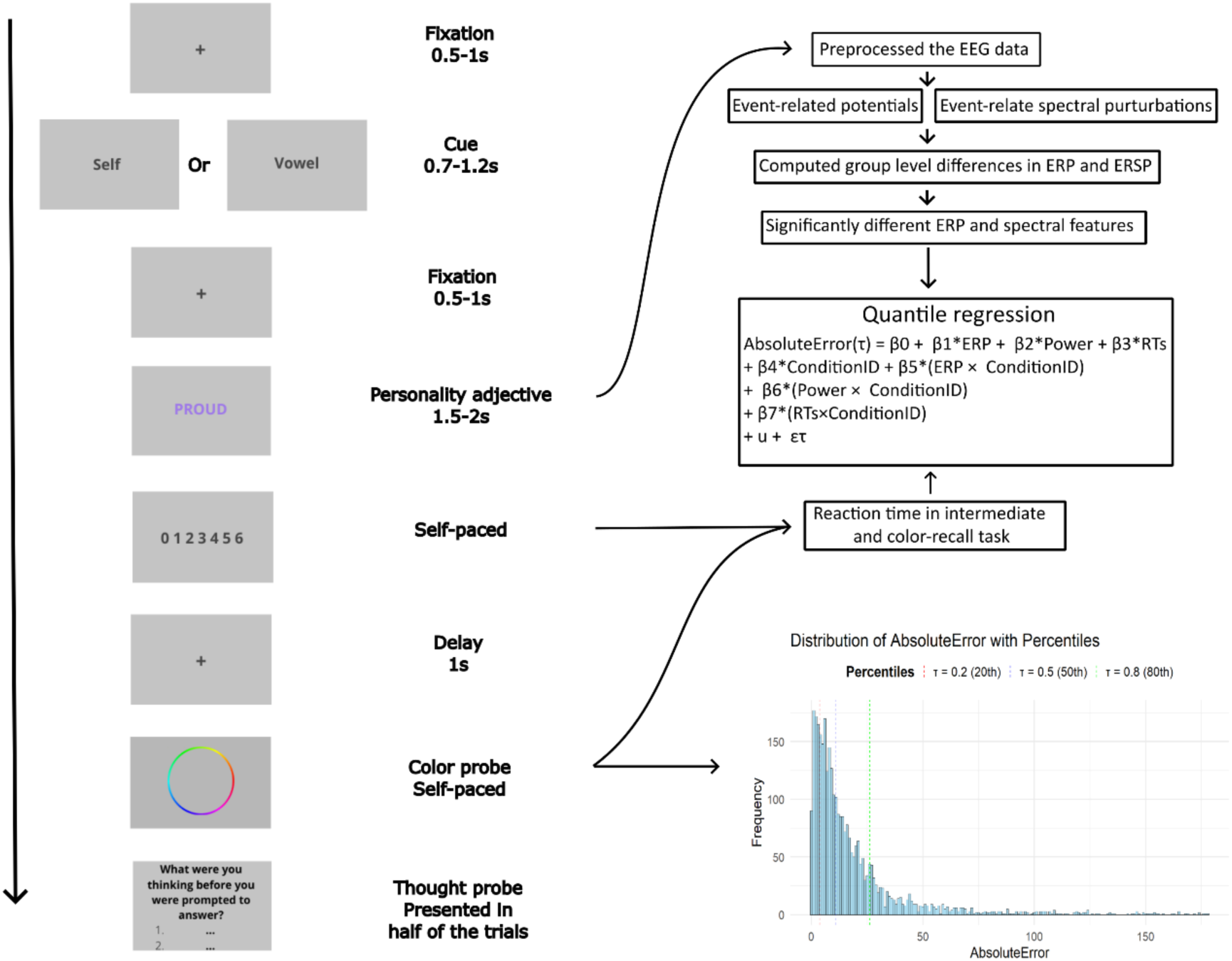
Behavioral paradigm and work-flow of the study. **Behavioral paradigm** (on the left)— Participants were given a cue, either ‘Self’ or ‘Vowel’ at the start of the trial. Then, a personality adjective was presented in a font color. The participant responded to the adjective according to the cue on a 7-point scale; if the cue was ‘self’, they respond to the question “How much this word describes your personality?”, and if the cue was ‘vowel’, they counted the number of vowels in the word. After a fixed-delay of 1 s, participant was probed for the memory of the font-color using a color wheel. In half of the trials randomly, a thought probe was presented at the end of the trial (see Methods section 2.2 for detail). **Work-flow of the study** (on the right)—EEG data was preprocessed and event-related potentials and spectral features were computed. From the behavioral data, reaction times and color-recall accuracy were computed. After a group-level analysis, significantly different EEG features between the two conditions, and reaction times were used as predictors in a conditional quantile regression model to estimate the color-recall performance at different quantiles (see Methods section 2.7 for details).

Following the cue, a fixation cross was displayed for a randomly varied duration between 0.5 s and 1 s. Next, a personality adjective (see **supplementary Table 1**) from a list of frequently used trait adjectives (Anderson, 1968) appeared in the centre of the screen, shown in a specific font color selected from hues of the color wheel (see section 2.3 for color-wheel design). The adjective was presented for a duration uniformly varied between 1.5 s to 2 s (mean = 1.78 s, SD = 0.16 s).

Depending on the cue, participants performed one of the two tasks:

- In EDA condition (vowel), participants counted the number of vowels in the word and responded on a seven-point scale indicating the count (0-6). To standardize response times and keep delays comparable across conditions (see **Supplementary Figure 1**), adjectives were selected to include between two and four vowels, based on pilot testing.
- In IDA condition (self), participants rated how well the adjective described their own personality on the seven-point scale, where 0 meant *“not at all me”* and 6 meant *“totally me”*.

After a fixed one second delay, a color wheel appeared on the screen to probe the font color of the adjective. Participants were instructed to respond as accurately and quickly as possible to both the rating and the color-recall task.

On 50% of the trials, selected randomly, a thought probe was presented after the color-recall response to assess the participant’s thought content during the delay period prior to color-recall (Huijser et al., 2018). The probe, adapted from prior studies (Huijser et al., 2018; Stawarczyk et al., 2011; Unsworth & Robison, 2016), asked: *“What were you thinking before you were prompted to answer?”*. Response options included: 1. *I tried to remember the color of the word*; 2. *I was still thinking about the word from the decision task*, 3. *I was evaluating aspects of the task*, 4. *I was distracted by the environment or my physical state*; 5. *I was daydreaming/thinking about task-unrelated matters*; 6. *I was not paying attention, but did not think about anything specific*.

### 2.3 Stimulus design and presentation

The experiment was designed in Matlab® (The MathWorks, Inc., Natick, MA), using the Psychophysics Toolbox (Brainard, 1997; Kleiner et al, 2007; Kleiner, 2007; Pelli, 1997) and displayed on a 22-inch LED monitor screen (75 Hz; 1440 x 900 pixels) at a viewing distance of approximately 75 cm. The screen subtended a visual angle of 35.14° (width) and 22.40° (height). The stimuli were presented on a gray background (128,128,128; RGB255), and all text was generated in 50pt. Arial font. The color wheel was designed in RGB color space with hue angles mapped directly on the spatial angles (a total of 360 hues from 0° to 360°, with each hue incremented by 1°, inner radius = 6.44°, outer radius = 7.85°). The hue angle and spatial angle were kept constant throughout the experiment, so that each degree on the wheel uniquely corresponded to a specific hue. For each trial, the font color of the personality adjective was randomly selected from these 360 evenly spaced hues on the wheel (e.g., a hue of angle 0° correspond to red, 120° to green, and 240° to blue; see Figure 1). Participants completed 120 trials in total, consisting 60 trials per condition. The experiment was divided into three blocks of 40 trials each. There was a separate practice block of 12 trials with a different personality adjective list.

### 2.4 EEG data acquisition

EEG and behavioral data were acquired in a sound-attenuated room, and the ambient light was kept the same in all the recording sessions. EEG was acquired at a sampling frequency of 1 kHz using a 64-channel ActiChamp (Brain Products, Germany) with active electrodes for a better signal-to-noise ratio. The electrode placement used the 10% electrode placement system. The impedance was maintained below 15kΩ and checked before and after the experiment. Electrode FCz was taken as a reference while recording the data. The Psychtoolbox was synchronized to the EEG acquisition system by sending triggers through parallel ports from the computer used to present stimuli to the EEG data acquisition computer.

### 2.5 Behavioral data analysis

Color recall accuracy was quantified by measuring the absolute angular error (absolute error), defined as the absolute difference between the presented hue and the hue reported by the participant. Reaction times for the intermediate task and color recall task were analysed. For thought-probe analysis, participant-wise percentage response of each option was calculated, and paired Wilcoxon signed rank test were conducted to compare percentage responses between the IDA and EDA conditions across the six options. Consistent with Huijser et al., 2018, the response option 1 was labelled as *on-task*, option 2 as *mental-elaboration*, option 3 as *task-related interference*, option 4 as *external distraction*, option 5 as *mind-wandering*, and option 6 as *inattentiveness*. All the responses, excluding *on-task* (option 1), were referred to jointly as *off-task*.

### 2.6 EEG data preprocessing and analysis

EEG data were analysed in MATLAB version R2023b (www.mathworks.com) using custom scripts as well as EEGLAB (Delorme & Makeig, 2004) and ERPLAB (Lopez-Calderon & Luck, 2014) toolbox functions. First, the DC offset was removed, and the data were high-pass filtered at 0.1 Hz using a non-causal Butterworth filter. Data segments corresponding to break periods between trial blocks were removed. Channels exhibiting excessive noise—defined as prolonged flatlines, high variance, or extreme amplitude fluctuations—were interpolated using spherical spline interpolation (Perrin et al., 1989; mean number of channels interpolated = 1.1).

To prepare the data for independent component analysis (ICA), segments exceeding a threshold of 500 µV within any 250 ms window (stepped every 50 ms) were removed. The data were then low-pass filtered at 45 Hz and down sampled to 250 Hz. ICA was performed using the infomax algorithm (*runica),* implemented in EEGLAB. Components were visually inspected, and those corresponding to ocular and muscular artifacts were removed (mean number of components removed = 9.6, SD = 4.7). The data were then re-referenced using the common-average re-referencing scheme.

The epochs corresponding to the encoding and delay periods were segmented for further analysis. The encoding epoch was defined from −500 ms to +1200 ms relative to the onset of the personality adjective stimulus. The delay epoch spanned a fixed 1-second period immediately preceding the onset of the color recall wheel (see **Figure 1**). Further, to reject commonly recorded artefactual potentials, which include skin potentials, movement artifacts, sudden voltage changes of unknown origin, the simple voltage threshold (SVT) and moving window peak-to-peak (MW) algorithm were used on the epoched data (Kappenman et al., 2021; Lopez-Calderon & Luck, 2014). A voltage threshold of −150 to 150 µV for SVT and a window size of 500 ms with a window step of 100 ms for MW was used. On average, 5.12% of the trials per participant were rejected.

Event-related potentials (ERP) were computed for the encoding epoch, and the pre-stimulus duration of 500 ms was used for baseline subtraction. Mean amplitude was computed for the P200 component (152–252 ms post-stimulus) and the late positive potential (LPP) (400–1196 ms) based on prior literature (Auerbach et al., 2016; Katyal et al., 2020; Schindler et al., 2023).

Time-frequency analysis was performed using a complex Morlet wavelet transform across the 3–40 Hz range. The number of wavelet cycles increased linearly from 2 (at 3 Hz) to 12.5 (at 40 Hz), allowing for an optimal trade-off between temporal and frequency resolution. The resulting wavelet temporal window ranged from approximately 666 ms at 3 Hz to 313ms at 40 Hz, power values were baseline-corrected by subtracting the average power in the pre-stimulus window (−400 to 0 ms) from post-stimulus power estimates. The FieldTrip toolbox’s (Oostenveld et al., 2011) permutation testing, based on t-statistics computed at each time-frequency point and combined with cluster correction for multiple comparisons, was used to estimate significantly different time-frequency clusters between the conditions (Maris & Oostenveld, 2007). 10000 permutations were computed, and the false positive (alpha) threshold was kept at 0.05 in all the statistical analyses.

### 2.7 Quantile regression

To study the relationship between EEG features, reaction times (RTs), and task-performance measured via recall error, conditional quantile regression was performed using *lqmm* package version 1.5.8 (Geraci, 2014) in R version 4.4.1 (http://www.R-project.org). Participants’ color recall error showed a right-skewed distribution, with many small errors and fewer large deviations (see **Figure 1**; bottom right). To account for this variability, the models were estimated at three quantiles:

- τ = 0.2 (lower quantile), capturing factors influencing small errors.
- τ = 0.5 (median quantile), reflecting the central tendency of errors.
- τ = 0.8 (upper quantile), capturing factors influencing large errors, when participants deviated substantially from the correct response.

To identify the optimal set of predictors for recall error, three hierarchical quantile regression models of increasing complexity were constructed and compared. The selection of predictors was informed by the group-level significant differences observed in the EEG features between the conditions (see Results section 3.2 and 3.3). Each trial was treated as an independent observation, with SubjectID included as a random intercept to account for inter-subject variability.

- Model 1: Behavioral predictors and their interactions with condition type.

This model included only the condition type *(ConditionID)*, reaction times from the intermediate *(RT_I_)* as well as the color-recall task *(RT_C_)*, and their interaction with condition type as predictors.

- Model 2: Behavioral predictors, EEG features from encoding epoch, and their interactions with condition type.

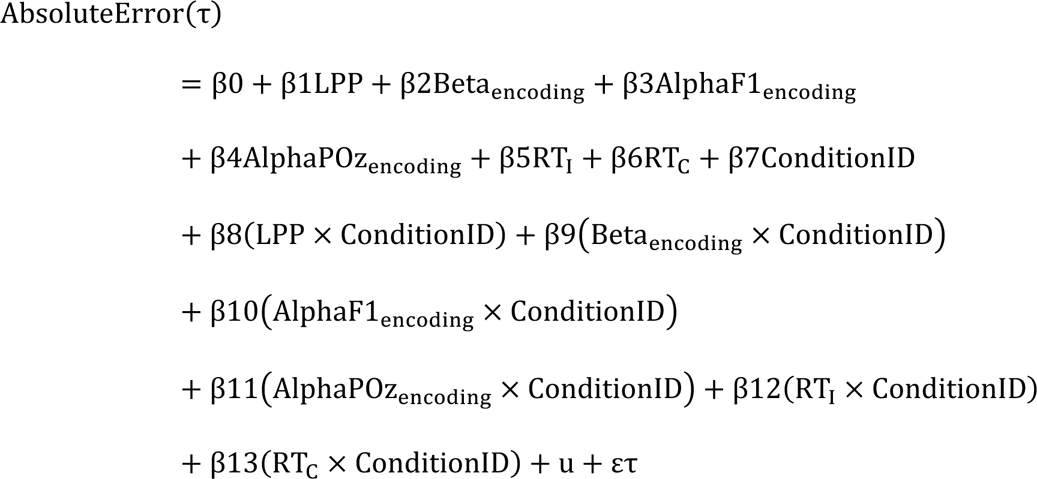

Here, EEG features quantified during the encoding (LPP amplitude, beta (15-25 Hz; averaged over 300 to 800 ms) and alpha (8-12 Hz; averaged over 600 to 800 ms) power over frontal electrodes, and alpha power (8-12 Hz; averaged over 252 to 800 ms) over parietal-oc (1) electrodes) were added as main effects along with their interactions with condition type. u is the random intercept for each subject and ετ represents the residuals at each quantile τ.

- Model 3: Behavioral predictors, EEG features from encoding epoch and delay epoch, and their interactions with condition type.

This final model, in addition to predictors from Model 1 and 2, further included EEG features of the delay period along with their interactions with condition type as main effects.

The model fit was compared using Akaike Information Criterion (AIC) and the model with the lowest AIC was selected as an optimal model and used for further analysis (**Table 1**). The *lqmm* package computed p-values using a block-bootstrap approach (Geraci, 2014; Geraci & Bottai, 2014), a robust method specifically chosen for its ability to handle clustered data in mixed models and to provide accurate inference without strong distributional assumptions. To account for multiple comparisons across predictors, the Benjamini-Hochberg False Discovery Rate (FDR) correction (Benjamini & Hochberg, 1995) was applied to the p-values. This approach controls the expected proportion of false positives while maintaining statistical power.

**Table 1.**
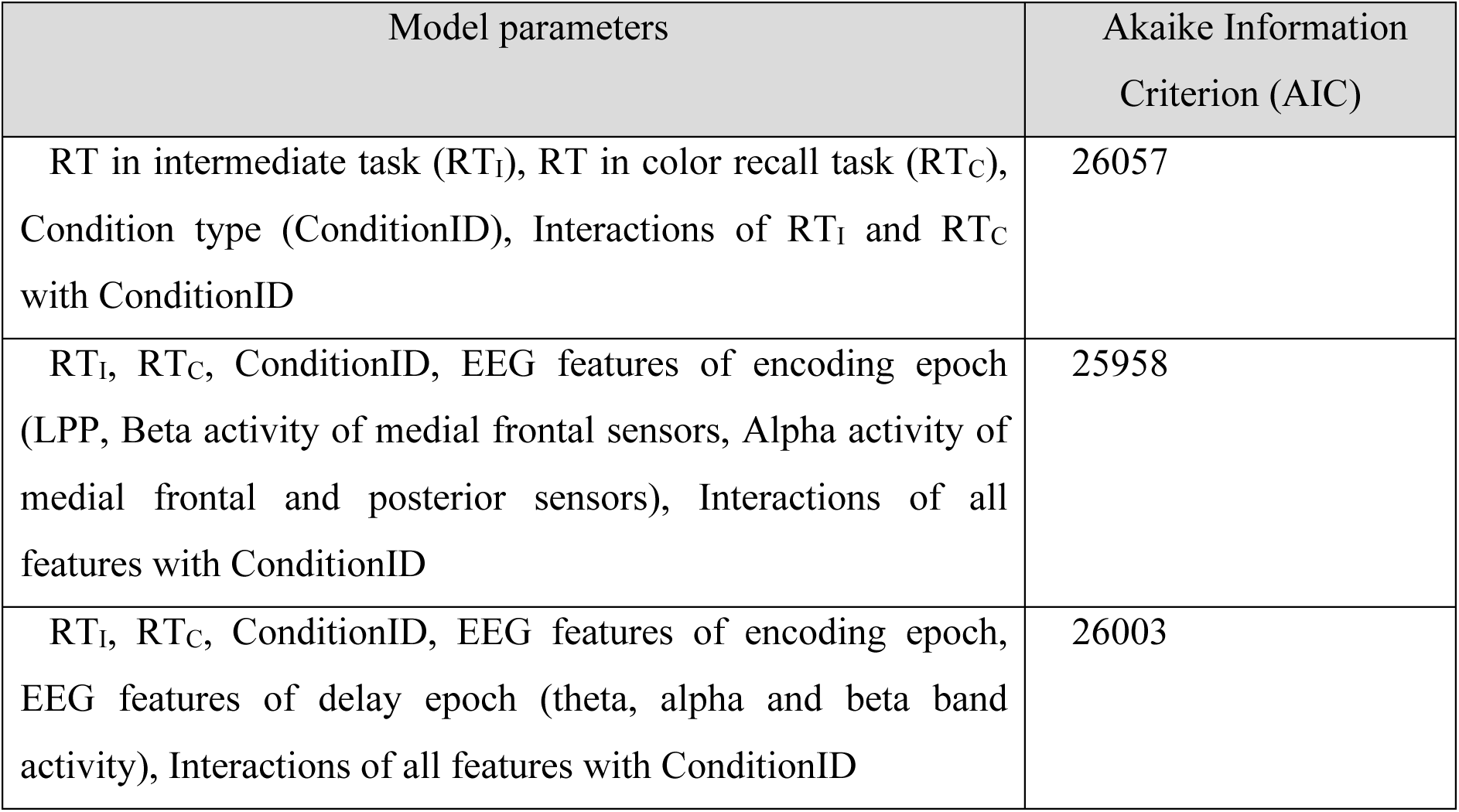
Model comparison based on Akaike Information Criterion (AIC) values. Lower AIC values indicate better fit. Model 1 included behavioral variables and their interactions with condition type. Model 2 added EEG features from the encoding epoch along with their interactions with condition type. Model 3, in addition to all the predictors of Model 2, added EEG features from the delay epoch.

| Model parameters | Akaike Information Criterion (AIC) |
| --- | --- |
| RT in intermediate task (RT<sub>I</sub>), RT in color recall task (RT<sub>C</sub>), Condition type (ConditionID), Interactions of RT<sub>I</sub> and RT<sub>C</sub> with ConditionID | 26057 |
| RT<sub>I</sub>, RT<sub>C</sub>, ConditionID, EEG features of encoding epoch (LPP, Beta activity of medial frontal sensors, Alpha activity of medial frontal and posterior sensors), Interactions of all features with ConditionID | 25958 |
| RT<sub>I</sub>, RT<sub>C</sub>, ConditionID, EEG features of encoding epoch, EEG features of delay epoch (theta, alpha and beta band activity), Interactions of all features with ConditionID | 26003 |

## Results

### 3.1 Participants reported more *off-task* in IDA

Participants’ responses to the thought-probe were analyzed to identify their brain state prior to the color-recall task. Since the thought probe required participants to report their mental states in six categories (see Methods section 2.2 for details), the percentage of responses to each option was subsequently calculated for each participant. Based on an earlier study (Huijser et al., 2018), different states of attention corresponding to each option was identified where option 1 was classified as *on-task*, option 2 as *mental elaboration*, option 3 as *task-related interference*, option 4 as *external distraction*, option 5 as *mind wandering*, and option 6 as *inattentiveness*. The distribution of responses between EDA and IDA for each option was then compared using paired non-parametric tests.

The percentage of responses to the option 2, which is characterized as *Mental elaboration,* were significantly higher in IDA (Wilcoxon signed rank test, Z = 3.40, p = 0.0007, Cohen’s d = 0.67), whereas percentage of responses to option 1, characterized as *on-task.* were lower in IDA relative to EDA (Wilcoxon signed rank test, Z = −3.33, p = 0.0009, Cohen’s d = −0.61) (**Figure 2**). The responses to *task-related interference, external distraction, inattentiveness, and mind wandering* were not significantly different between the IDA and EDA condition.

**Figure 2.**
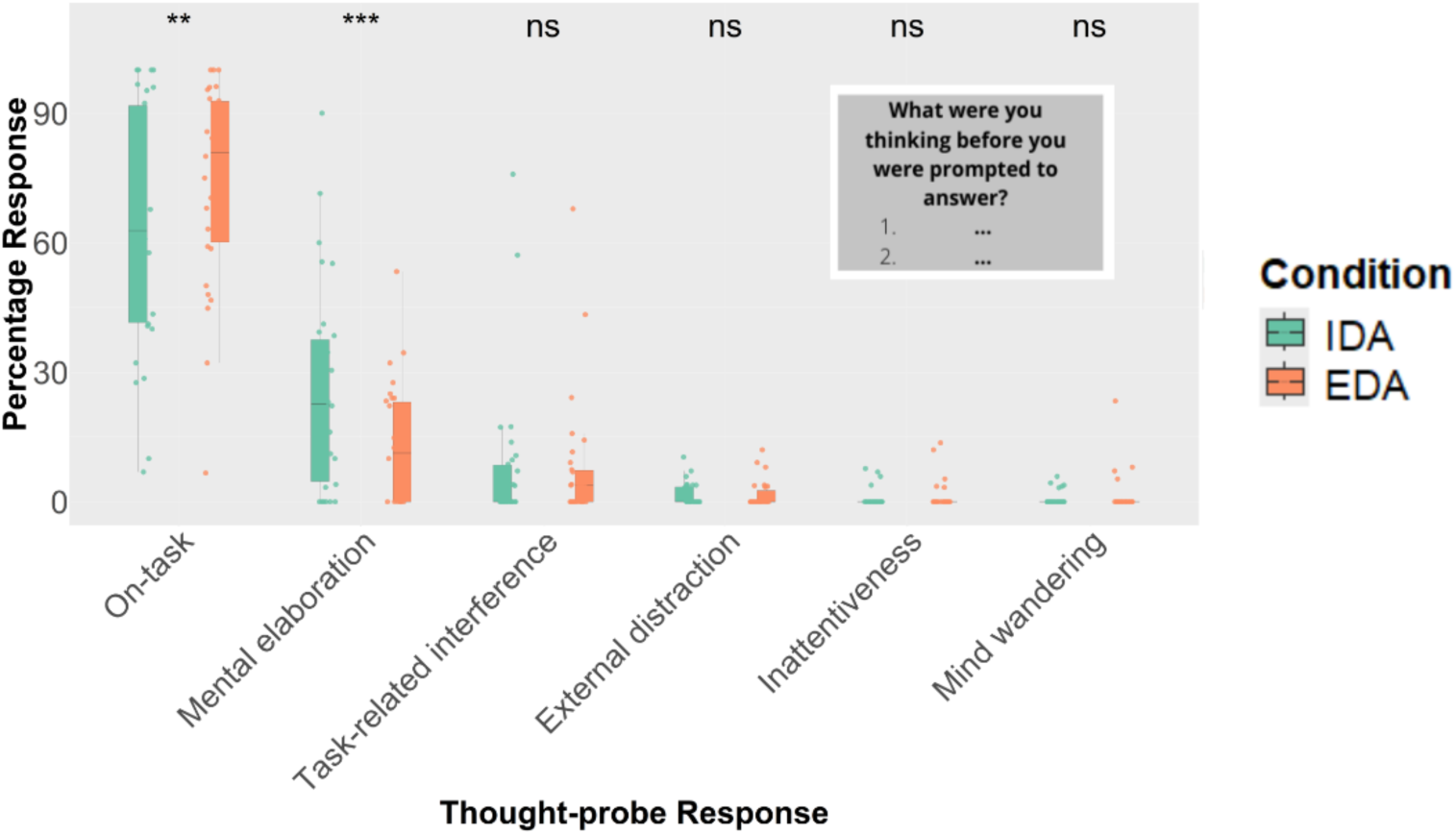
Response in the thought-probe plotted as percentage response for each option of the thought probe. Higher frequency of mental elaboration of the word and lower frequency of on-task in IDA relative to EDA.

This self-report indicates a higher frequency on mental-elaboration of personality adjective during the delay period in IDA condition, supporting the validity of using the cue to manipulate the processing of word across conditions. Further this higher frequency of mental-elaboration could potentially hinder the rehearsal of the color to be recalled in the subsequent task, thereby impairing the performance in color-recall task.

### 3.2 Higher amplitude of late-positive potentials in IDA

The scalp distribution of the instantaneous voltages was visualized at time points ranging from 0 ms to 1000 ms, in 100 ms steps, following the onset of the word. This captured the temporal evolution and the sustained processing of the personality adjective over the scalp (see **supplementary Figure 2**). These scalp maps showed that, in IDA condition, processing was predominantly sustained over frontal electrodes, whereas in the EDA condition, it was more pronounced over parietal electrodes. This observation informed the decision to focus on frontal and parietal electrodes in the subsequent analyses.

For P200 component, no significant difference between IDA and EDA was observed at frontal electrodes (Fz; t (27) = −1.01, p = 0.32, Cohen’s d = −0.19). Late positive potentials (LPP) over frontal electrodes were significantly higher in amplitude in IDA compared to EDA (Fz; t (27) = 5.16, p < 0.0001, Cohen’s d = 0.97) (**Figure 3a**). Over parietal electrodes, the LPP between conditions was not significantly different (t (27) = −0.26, p = 0.64, Cohen’s d = −0.08). LPP are linked to affective processing of the stimuli (De Cesarei & Codispoti, 2011; Dennis & Hajcak, 2009; Naumann et al., 1992), suggesting that processing the word in the IDA condition led to affective engagement compared to counting the vowels in the EDA condition.

**Figure 3.**
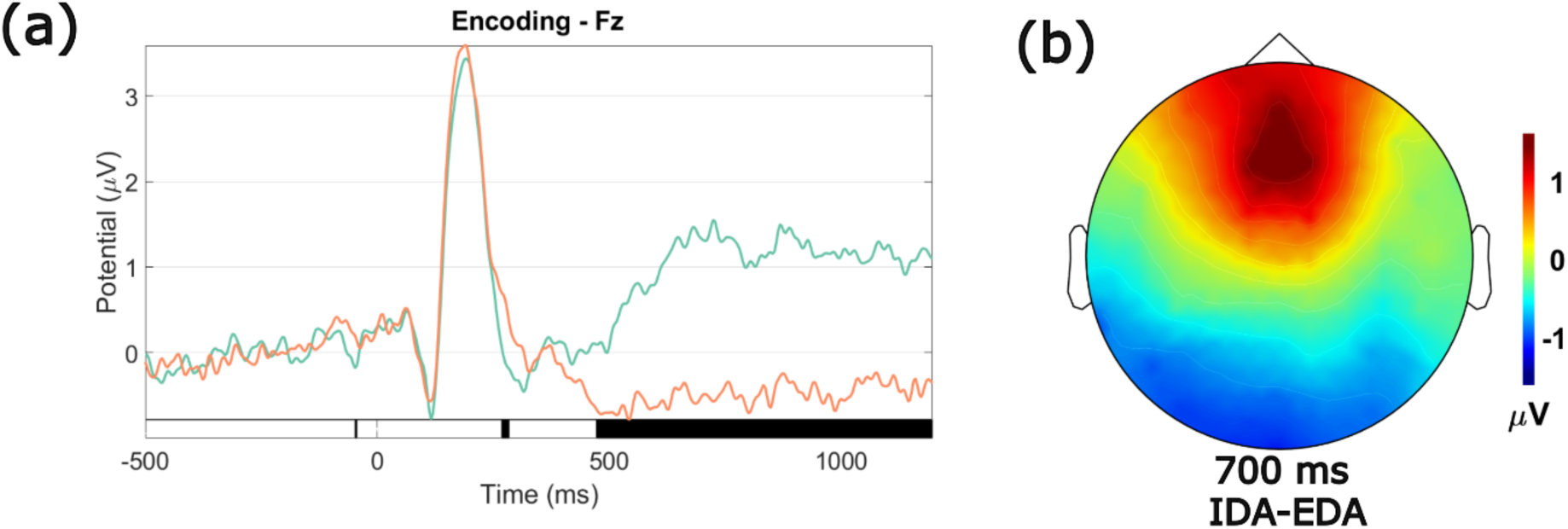
Late-positive potentials (LPP) in encoding the word in IDA. (a) Event related potential, during encoding of the word, at medial-frontal sensor (Fz for representation). Higher amplitude in late-positive potentials in IDA. Black bar below at the bottom plots statistically significant differences (alpha threshold at 0.05). (b) Topo-plot showing difference map (IDA minus EDA) of instantaneous voltage at 700 ms.

### 3.3 Spectral differences in IDA relative to EDA

Next, time-frequency analysis was performed on the EEG data to examine differences in spectral power between the two conditions. Specifically, spectral features were analyzed during two time-windows: the encoding period and the delay period immediately preceding the color-recall task. A complex Morlet wavelet transform was applied to perform time-frequency analysis across frequencies 3 to 40 Hz.

In the encoding epoch, a cluster-based permutation test based on t-statistics indicated a significant effect of condition (p < 0.05). In the observed data, this corresponded to a cluster showing higher event-related desynchronization (ERD) in beta frequency band activity in IDA compared to EDA over the medial-frontal sensors (F1, F2; see **Figure 4a).** Further, a significantly higher event-related desynchronization (ERD) was observed in the alpha frequency band over parieto-occipital sensors (POz; see **Figure 4b**) and medial-frontal sensors (F1, F2) in the IDA condition (see **Supplementary Figure 3** for time-frequency plots of each condition).

**Figure 4.**
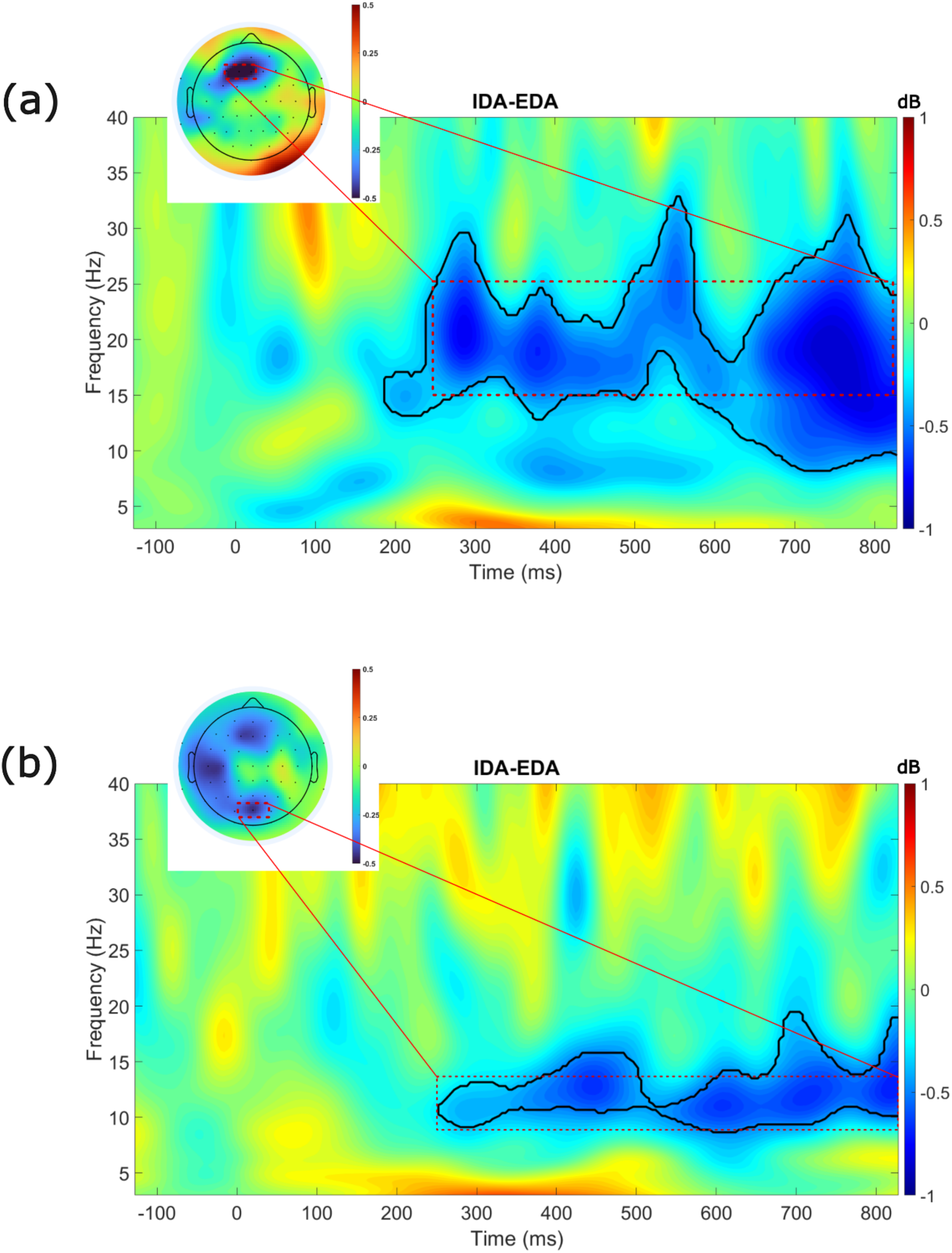
Time-frequency plots showing difference in power during encoding of the stimulus (IDA minus EDA). (a) The difference in power between IDA and EDA over medial frontal sensors (F1 plotted for representation) and (b) posterior sensors (POz plotted for representation). Insets show topographical distribution of the condition difference with electrode locations marked. Black contours outline cluster of significant difference (p < 0.05, corrected).

Additionally, during the delay period prior to the recall of the color, a higher alpha power was observed in IDA over occipito-parietal sensors (Oz, PO7, P7, P5, CP5, CP3) relative to the EDA condition (**Figure 5**).

**Figure 5.**
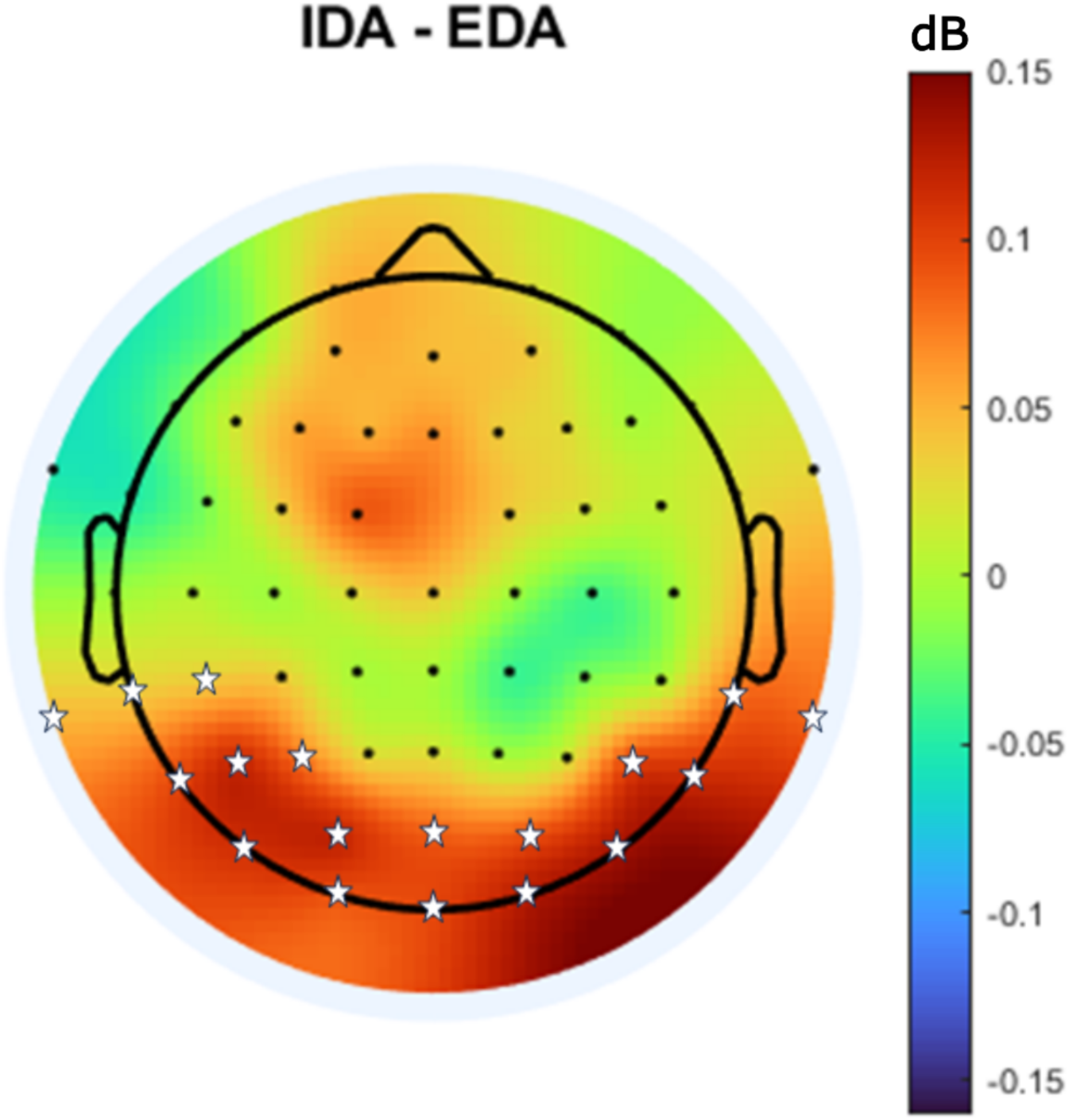
Topography plot of difference in in power between IDA and EDA in alpha band (8-12 Hz) during the delay period prior to the color-recall. Stars indicate statistically significant sensors (p < 0.05).

### 3.4 Beta power over medial-frontal sensors during encoding is a significant modulator in color-recall performance

To investigate how neural and behavioral predictors influence absolute error across different levels of performance, linear quantile mixed models (LQMM) were fitted at three quantiles: τ = 0.2 (lower errors), τ = 0.5 (median errors), and τ = 0.8 (higher errors) (see Methods section 2.7 for details). To identify the optimal model for predicting absolute error, a series of quantile regression models of increasing complexity were compared using AIC (**Table 1**). First, simpler models were tested with only RTs or condition (IDA vs. EDA) as predictors, followed by models that also included EEG features from the encoding and delay periods. Based on lowest AIC, the final model included LPP amplitude, medial-frontal beta and alpha during encoding, parietal-occipital alpha during encoding, RTs from the intermediate and color-recall tasks, and ConditionID as fixed effects. It also incorporated interactions between each predictor and ConditionID, as well as a random intercept for SubjectID (see Model 2 in Methods section 2.7). **Table 2** summarizes the parameter estimates, standard errors (SE), and p-values for each predictor at each quantile.

**Table 2.** Quantile regression results across three quantile levels (τ = 0.2, 0.5, 0.8). The models include behavioral predictors (reaction times from intermediate [RT_I_] and color-recall [RT_C_] tasks), EEG features from the encoding epoch (LPP amplitude, beta and alpha power at medial-frontal and posterior electrodes), condition type (ConditionID), and interaction terms between ConditionID and all predictors. Coefficient estimates, standard errors, and FDR-corrected p-values are reported. Shaded rows are statistically significant.

| Tau | Coefficient | Estimate | Std. Error | Adjusted P value<br>(FDR corrected) |
| --- | --- | --- | --- | --- |
| 0.2 | (Intercept) | 6.46011 | 1.433612 | 0.000431271 |
|  | LPP | 0.018589 | 0.036318 | 0.733282797 |
|  | Beta<sub>encoding</sub> | 0.134662 | 0.060314 | 0.079202359 |
|  | AlphaF1<sub>encoding</sub> | 0.017063 | 0.040083 | 0.742955958 |
|  | AlphaPOZ<sub>encoding</sub> | -0.03708 | 0.04676 | 0.598098771 |
|  | RT<sub>I</sub> | -0.02124 | 0.222588 | 0.946930883 |
|  | RT<sub>C</sub> | -1.07577 | 0.455886 | 0.062484773 |
|  | **ConditionID** | **-6.59793** | **1.849805** | **0.003126915** |
|  | LPP:ConditionID | 0.020955 | 0.04734 | 0.742955958 |
|  | **Beta<sub>encoding</sub>:ConditionID** | **-0.26149** | **0.083037** | **0.009010833** |
|  | AlphaF1<sub>encoding</sub>:ConditionID | 0.007239 | 0.051813 | 0.933932266 |
|  | AlphaPOZ<sub>encoding</sub>:ConditionID | -0.03667 | 0.061762 | 0.686126918 |
|  | RT<sub>I</sub>:ConditionID | 0.610178 | 0.278053 | 0.081460492 |
|  | **RT<sub>C</sub>:ConditionID** | **2.545962** | **0.69796** | **0.002690095** |
| 0.5 | (Intercept) | 9.239357 | 1.964283 | 0.00029678 |
|  | LPP | 0.042254 | 0.058023 | 0.598098771 |
|  | Beta<sub>encoding</sub> | 0.123628 | 0.120862 | 0.503006735 |
|  | AlphaF1<sub>encoding</sub> | 0.070542 | 0.09453 | 0.598098771 |
|  | AlphaPOZ<sub>encoding</sub> | -0.08548 | 0.08059 | 0.503006735 |
|  | RT<sub>I</sub> | 0.634443 | 0.340384 | 0.151051499 |
|  | RT<sub>C</sub> | 0.822708 | 1.107281 | 0.598098771 |
|  | **ConditionID** | **-9.10718** | **2.057913** | **0.000450896** |
|  | LPP:ConditionID | -0.07459 | 0.085946 | 0.564372257 |
|  | **Beta<sub>encoding</sub>:ConditionID** | **-0.47021** | **0.145172** | **0.007547003** |
|  | AlphaF1<sub>encoding</sub>:ConditionID | -0.05979 | 0.134328 | 0.742955958 |
|  | AlphaPOZ<sub>encoding</sub>:ConditionID | -0.01936 | 0.116086 | 0.933932266 |
|  | RT<sub>I</sub>:ConditionID | 0.558312 | 0.426296 | 0.377614059 |
|  | **RT<sub>C</sub>:ConditionID** | **4.362284** | **1.132003** | **0.001789954** |
| 0.8 | (Intercept) | 10.50498 | 2.076886 | 0.00013309 |
|  | LPP | -0.13649 | 0.085593 | 0.246147919 |
|  | Beta<sub>encoding</sub> | 0.004188 | 0.207683 | 0.983993724 |
|  | AlphaF1<sub>encoding</sub> | -0.10779 | 0.104387 | 0.503006735 |
|  | AlphaPOZ<sub>encoding</sub> | -0.16039 | 0.122849 | 0.377614059 |
|  | **RT<sub>I</sub>** | **1.892061** | **0.491238** | **0.001789954** |
|  | **RT<sub>C</sub>** | **5.490533** | **1.492325** | **0.002690095** |
|  | **ConditionID** | **-9.9743** | **2.314891** | **0.000551573** |
|  | LPP:ConditionID | 0.286378 | 0.119802 | 0.062150312 |
|  | Beta<sub>encoding</sub>:ConditionID | -0.30886 | 0.246571 | 0.394948654 |
|  | AlphaF1<sub>encoding</sub>:ConditionID | 0.214101 | 0.113076 | 0.149830178 |
|  | AlphaPOZ<sub>encoding</sub>:ConditionID | -0.16816 | 0.186256 | 0.556530437 |
|  | RT<sub>I</sub>:ConditionID | 0.479738 | 0.489292 | 0.515929338 |
|  | **RT<sub>C</sub>:ConditionID** | **6.152359** | **1.171858** | **0.00013309** |

Fixed effect of ConditionID revealed that participants made fewer errors in EDA relative to the IDA condition across all quantiles of the error distribution (**Figure 6a**). Interestingly, this effect becomes much stronger from lower quantiles (τ = 0.2, β = −6.59, SE = 1.84, p = 0.003) to median and upper quantiles (τ = 0.5, β = −9.10, SE = 2.05, p = 0.0004; τ = 0.8, β = −9.97, SE = 2.31, p = 0.0005). Reaction times did not have any meaningful impact on color recall in lower quantile i.e. τ = 0.2 (RT_I_, β = −0.02, SE = 0.22, p = 0.94; RT_C_, β = −1.07, SE = 0.45, p = 0.06) and median quantile i.e. τ = 0.5 (RT_I_, β = 0.63, SE = 0.34, p = 0.15; RT_C_, β = 0.82, SE = 1.10, p = 0.59) trials. However, slower responses in both the intermediate task and color recall task (RT_I_, β = 1.89, SE = 0.49, p = 0.001; RT_C_, β = 5.49, SE = 1.49, p = 0.002) contribute to higher errors (τ = 0.8) irrespective of the conditions (**Figure 6c** and **6d**). Interaction effects of medial-frontal beta power and ConditionID revealed that as beta power during encoding of the stimuli increased, error in color-recall decrease more sharply in EDA at τ = 0.2 (Beta_encoding × ConditionID; β = −0.26, SE = 0.08, p = 0.009) and τ = 0.5 (Beta_encoding × ConditionID; β = −0.47, SE = 0.14, p = 0.007; see **Figure 6b**), whereas no significant effects were observed at the upper quantile (τ = 0.8). Other EEG measures, including LPP amplitude and alpha power, did not meaningfully affect the color recall (see **Table 2**). Furthermore, a significant interaction between color-recall RT and ConditionID was present across all quantiles; specifically, in the EDA condition, longer color-recall RTs were linked to greater errors (τ = 0.2: β = 2.54, SE = 0.69, p = 0.002; τ = 0.5: β = 4.36, SE = 1.13, p = 0.001; τ = 0.8: β = 6.15, SE = 1.17, p = 0.0001), suggesting that reaction time effects are more pronounced when color-recall errors are large.

**Figure 6.**
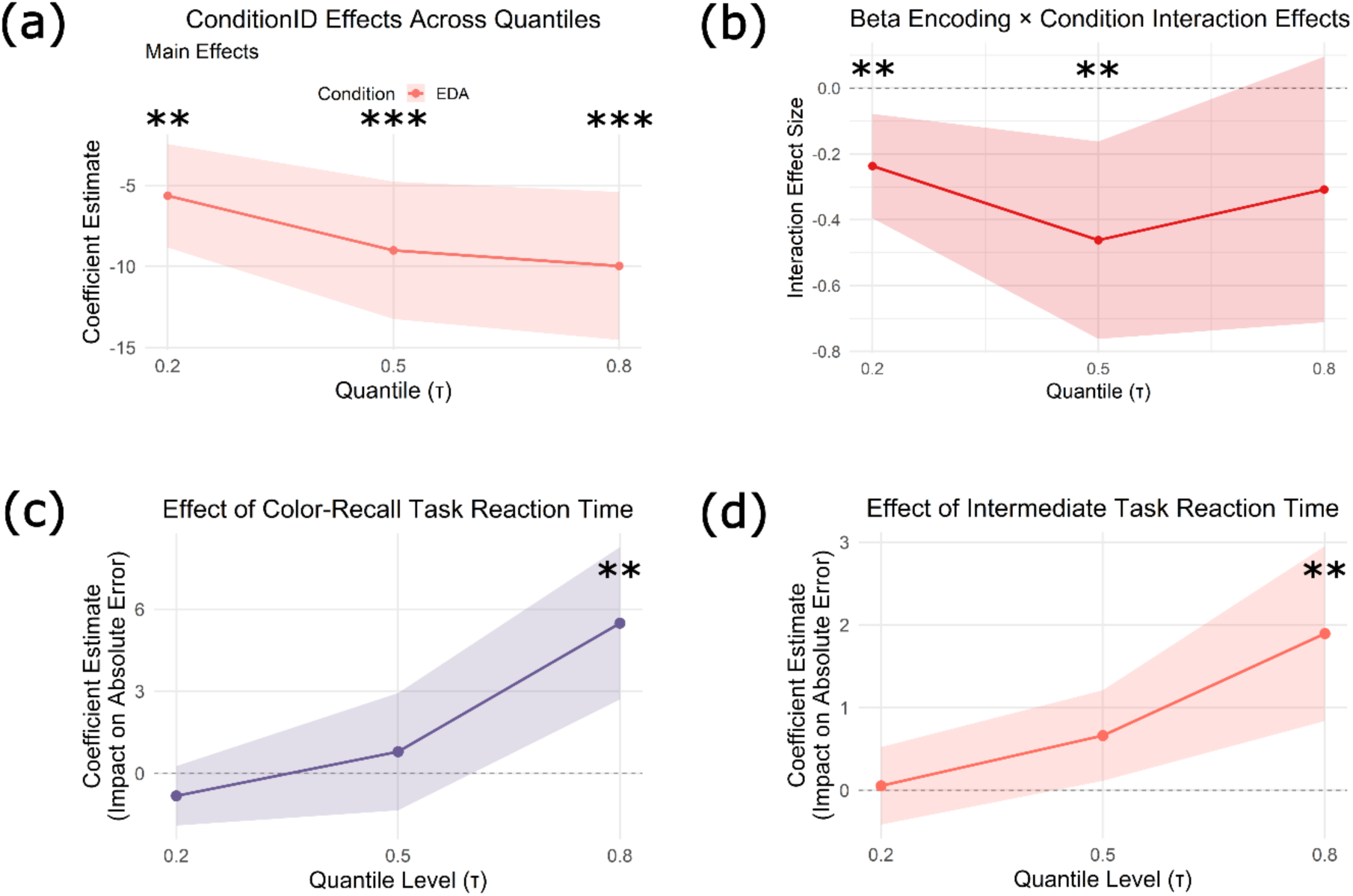
Effect of neural and behavioral predictors across quantiles of error response. (a) Estimated coefficients for the main effect of ConditionID (EDA) across quantiles. (b) Interaction effect between beta power during encoding and condition (EDA vs. IDA) across quantiles. (c) Main effect of reaction time in color recall task on error in color recall. (d) Main effect of reaction time in intermediate task on color recall. The shaded area represents the 95% confidence interval. Asterisks denote statistically significant effect (FDR corrected).

## Discussion

This study investigated the effects of internally directed attention (IDA) on working memory and the underlying neural mechanisms associated with such effects. A self-referential processing task was employed with a color recall working memory paradigm. In contrast, for externally directed attention (EDA) condition, a vowel-counting task was used instead of the self-referential task. The main empirical findings of the study were as follows, *First*, behaviorally, it was found that participants had higher accuracy in color recall in EDA condition compared to the IDA condition across all levels of performance. Additionally, participants’ color recall error response followed a right-skewed distribution, with many small errors and fewer large errors. *Second*, late positive potentials (LPP) were observed over medial frontal sensors during stimulus encoding in the IDA condition, suggesting affective processing of the personality adjectives. Additionally, event-related desynchronization (ERD) in the alpha and beta bands over medial-frontal sensors, and in alpha band over posterior sensors was greater in IDA relative to the EDA condition during encoding. During the delay period, higher alpha power was observed in IDA over occipito-parietal sensors compared to the EDA condition. *Third*, medial-frontal beta power in encoding, in interaction with condition type, significantly influenced color-recall accuracy at low and median quantiles of error distribution, whereas responses in high quantiles were significantly driven by reaction times.

Prior studies have shown that self-referential processing triggers self-generated thoughts (Huijser et al., 2018), which are considered a core component of IDA (Dixon et al., 2014). This paradigm, therefore, provides a robust approach to examining IDA mechanisms. The present thought-probe analyses revealed that participants were more off-task during the delay period in the IDA condition (**Figure 2**), evidenced by a higher frequency of mental elaboration. Consistent with these findings, color-recall performance was significantly poorer in IDA trials compared to EDA trials. These results suggest that self-referential processing may divert cognitive resources from the rehearsal of perceptual features, thereby impairing task performance.

Electrophysiological analyses confirmed that the personality adjectives were processed differently across conditions. Previous research on self-referential processing using EEG has shown distinct ERP signatures reporting the higher amplitude of LPP in response to personality adjectives (Katyal et al., 2020). In the present paradigm as well, processing the personality adjective in self-reference, as cued at the beginning of the trial, led to higher amplitude in LPP in relation to counting the number of vowels in the word (**Figure 3a**). LPPs are linked to sustained emotional engagement and affective processing of the stimulus and are modulated by the emotional relevance of the stimulus (Dennis & Hajcak, 2009; Hajcak et al., 2009; Lang & Bradley, 2010; Naumann et al., 1992; Schindler et al., 2023; Schupp et al., 2000). The P200 component in ERP, which is reported to be responsible for exogenous attentional capture (Carretié, 2014; Kanske et al., 2011) and automatic semantic processing of the word (Crowley & Colrain, 2004; Lei et al., 2017; Schindler et al., 2023), was not different between conditions suggesting that the early-stage processing might not differ between conditions. Further, the observed differences in LPP components during the processing of the personality adjective align with the time-course model proposed by Grainger & Holcomb (2009), which suggests that semantic elaboration and integration processes occur after 400 ms of stimulus presentation, corresponding to the time window in which the LPP differences were observed in this study. Thus, taking the current electrophysiological results and the literature into account, one can concur that the personality adjective is processed affectively in IDA compared to the EDA condition.

Time-frequency analyses revealed a greater alpha desynchronization during the encoding of personality adjectives in the IDA condition over both medial-frontal and parietal sensors (**Figure 4)**. These observations further strengthen the inference in the literature that alpha desynchronization correlates with emotional engagement with the stimuli (De Cesarei & Codispoti, 2011; Knyazev et al., 2008; Otten & Jonas, 2014; Schubring & Schupp, 2019). Conversely, in the delay period prior to color-recall, higher alpha power over posterior sensors was observed in IDA (**Figure 5**). This alpha increase in the delay period is a signature of internally directed attention (Kam et al., 2018; O’Connell et al., 2009). Taking together, the thought-probe results, ERP analyses, and the time-frequency findings indicate that the IDA condition promotes both affective processing during stimulus encoding and internal focus during the delay period, making this paradigm a robust choice to study IDA mechanisms. However, one caveat of the present design is that, although IDA is examined in contrast to EDA, certain cognitive systems such as those involved in goal maintenance and working memory are engaged in both conditions, making it challenging to unequivocally isolate neural processes purely specific to IDA.

Quantile regression was employed to assess the differential contributions of ERP, beta, and alpha power, and reaction times under internal and externally directed attention conditions. This is because the traditional regression methods, such as ordinary least squares (OLS), estimate the mean effect of the predictors on an outcome variable assuming a constant relationship across all levels of the dependent variable. However, in present study, given that the distribution of color-recall error is not homogenous across its range, the quantile regression examined how the independent variables predict different parts of the error distribution, thus allowing us to investigate effects at different levels of task performance (lower, median, and high errors in color-recall, corresponding to lower, median and upper quantile respectively). The effect of condition type (IDA vs. EDA) was significant across all quantiles, indicating a consistent performance advantage in the EDA condition. However, the magnitude of this effect increased at median and upper quantiles, suggesting that while the condition difference was already present in low-error trials, it became even more pronounced in trials with higher error in color-recall. While group-level differences in alpha and LPP measures were evident between the conditions, they did not significantly predict performance at the single-trial level. A possible explanation could be that these measures reflect neural processes that are not directly linked to performance in the color-recall task in the current paradigm but are just the signatures of internally directed attention.

The IDA condition likely requires deeper semantic and self-referential processing. This leads to more self-generated thoughts in IDA compared to EDA. The activation of medial prefrontal cortex (mPFC) is reported in both generation and retrieval of self-generated information (Subramaniam et al., 2012, 2019; Vinogradov et al., 2008). An MEG study by Subramaniam et al., (2019) further reported the suppression of mPFC beta power during self-generated information. In line with the previous findings, a reduction of beta power over medial-frontal sensors in IDA relative to EDA in the encoding epoch was observed. Further, it was observed that the extent of beta power change at frontal sensors had a significant impact on color-recall. Thus, more self-generated thoughts in IDA, as evidenced by thought-probe (**Figure 2**) and neural signatures (**Figure 4**), could lead to poor performance on color-recall. This can, in turn, happen because of two different mechanisms. More self-generated thoughts during the encoding phase of the trial in IDA could lead to cortical information processing facilitated by alpha and beta desynchronization (Hanslmayr et al., 2012). However, this reduces the resources for encoding the low-priority (font color) perceptual features and thus affects their recall. The alternative mechanism that could be at play is during the delay epoch prior to color recall where, internally directed attention in the delay period, as indexed by a concomitant increase in alpha activity (**Figure 5**) (Kam et al., 2018), could lead to mental elaboration of the word from the intermediate task (**Figure 2**) (Huijser et al., 2018). This leads to poor rehearsal of font-color and thus poor performance in the color-recall task. Since the time-frequency parameters of the delay period did not improve the quantile regression model fit, and the medial-frontal beta power during encoding did influence the performance at a single-trial level (**Figure 6b**), the current study provides evidence in support of the former mechanism.

In conclusion, the predominance of low error trials (see **Figure 1**; bottom right) indicate that the participants performed relatively well on the color recall task. Consequently, the quantile regression analyses reveal that medial-frontal beta power during encoding significantly predicts performance at these quantiles. Conversely, reaction times became dominant predictors only in the high-error trials, which are relatively few, indicating a shift to speed-accuracy tradeoff in such trials. Moreover, by integrating trial-level EEG with quantile-regression, we demonstrate that IDA diverts resources during encoding, suggesting impaired perceptual binding. The observed distinct oscillatory mechanisms (alpha/beta) across task phases predict how IDA impacts visual working memory processing during goal-directed tasks.

## Data and Code Availability

*The raw EEG data used in this study is available from the corresponding authors upon reasonable request. However, the processed EEG data and all the relevant codes used for subsequent analysis in this study are available in the provided link* https://drive.google.com/drive/folders/19X5xdNLVueJf5NniAL4JtUKgXewgqWEH

## Declaration of Competing Interest

The authors declare no competing interest that could influence the work reported in this article.

## Author Contributions

**Ankit Yadav:** Conceptualization, Experimental design, Data collection, Formal analysis, Methodology, Visualization, Writing – Original draft preparation. **Arpan Banerjee:** Funding acquisition, Project administration, Resources, Supervision, Validation, Writing – Review & editing. **Dipanjan Roy:** Conceptualization, Experimental design, Funding acquisition, Methodology, Project administration, Resources, Supervision, Validation, Writing – Review & editing.

## Acknowledgements

We thank Vinsea Singh for her helpful comments on improving the readability of the manuscript. NBRC Core funds supported this study. DR was supported by SERB Core Research Grant (CRG) S/SERB/DPR/20230033 extramural grant from the Department of Science and Technology, Ministry of Science and Technology, Govt. India. DR and AB acknowledge the generous support of the NBRC Flagship program BT/MIDI/NBRC/Flagship/Program/2019: Comparative mapping of common mental disorders (CMD) over the lifespan.

---

**Supplementary Figure 1.**
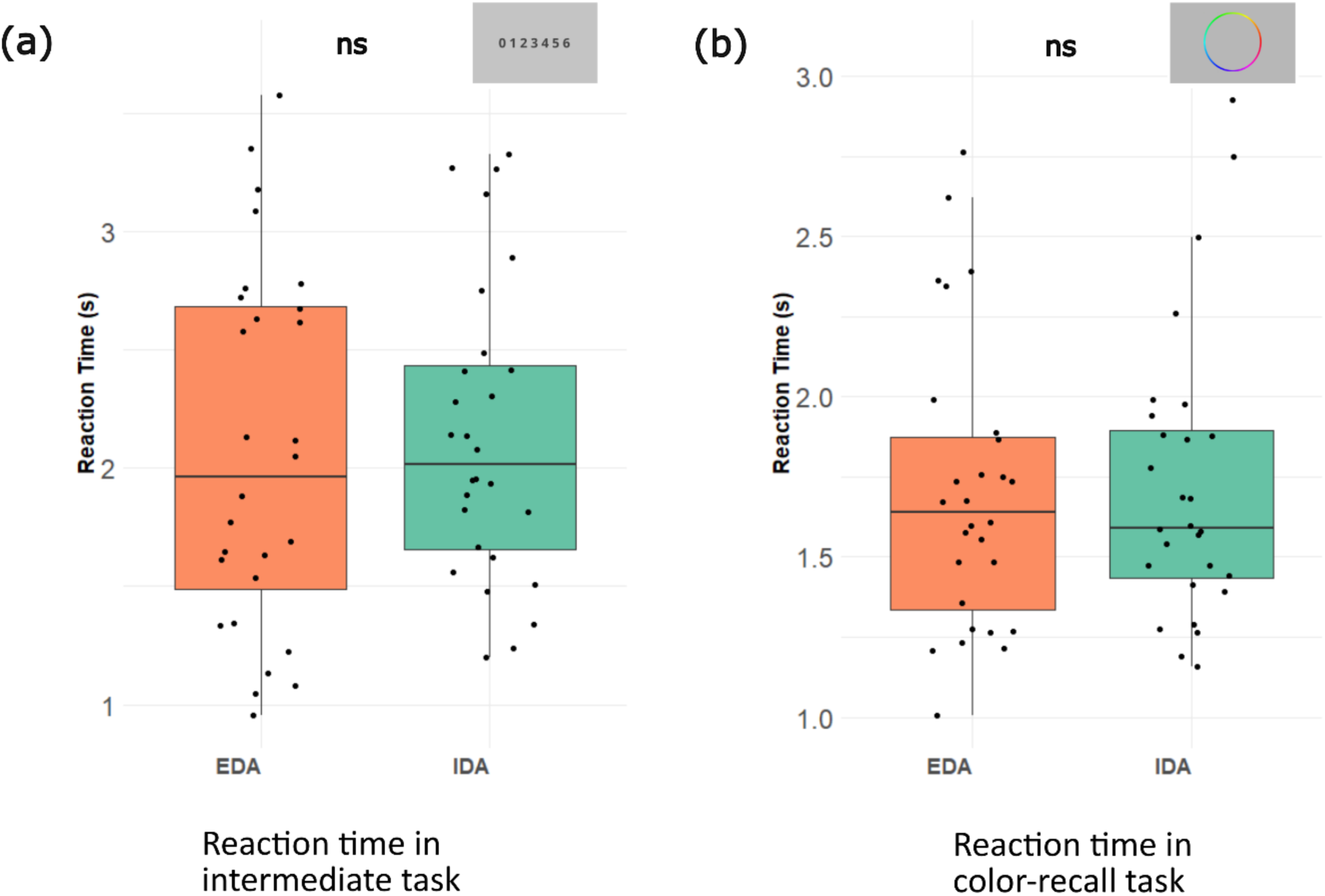
Reaction times in intermediate task and color-recall task. Personality adjectives were restricted to words containing two to four vowels in EDA condition, based on pilot testing to equalize reaction times across conditions and maintain a consistent overall delay duration. (a) Reaction time in intermediate task. (b) Reaction time in color-recall task.

**Supplementary Figure 2.**
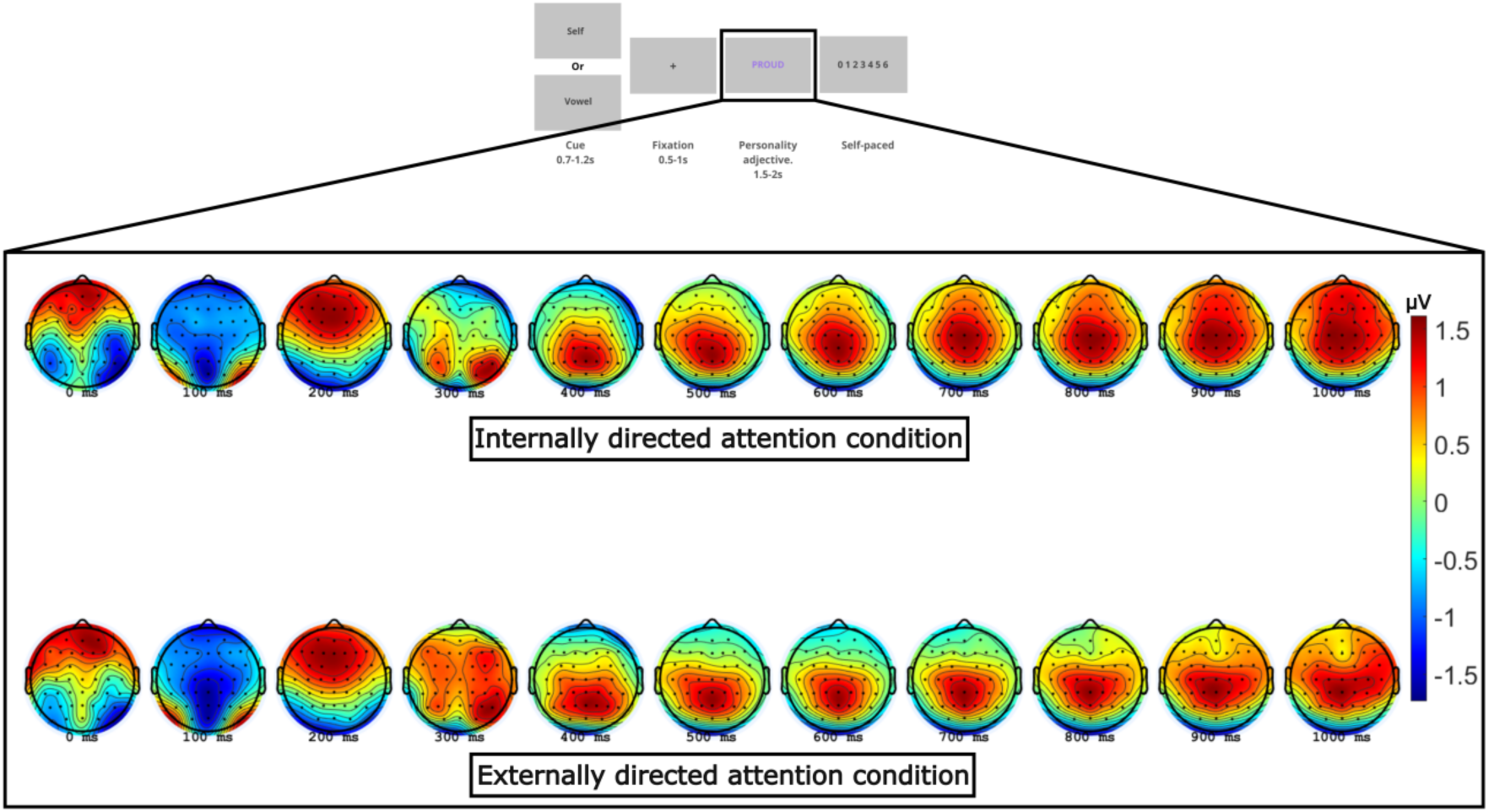
Scalp distribution of instantaneous voltage during the encoding epoch. The top row shows voltage distribution during self-referential processing of the personality adjective, while the bottom row shows voltage distribution during vowel-counting of the adjectives. Voltages are averaged over trials and across participants.

**Supplementary Figure 3.**
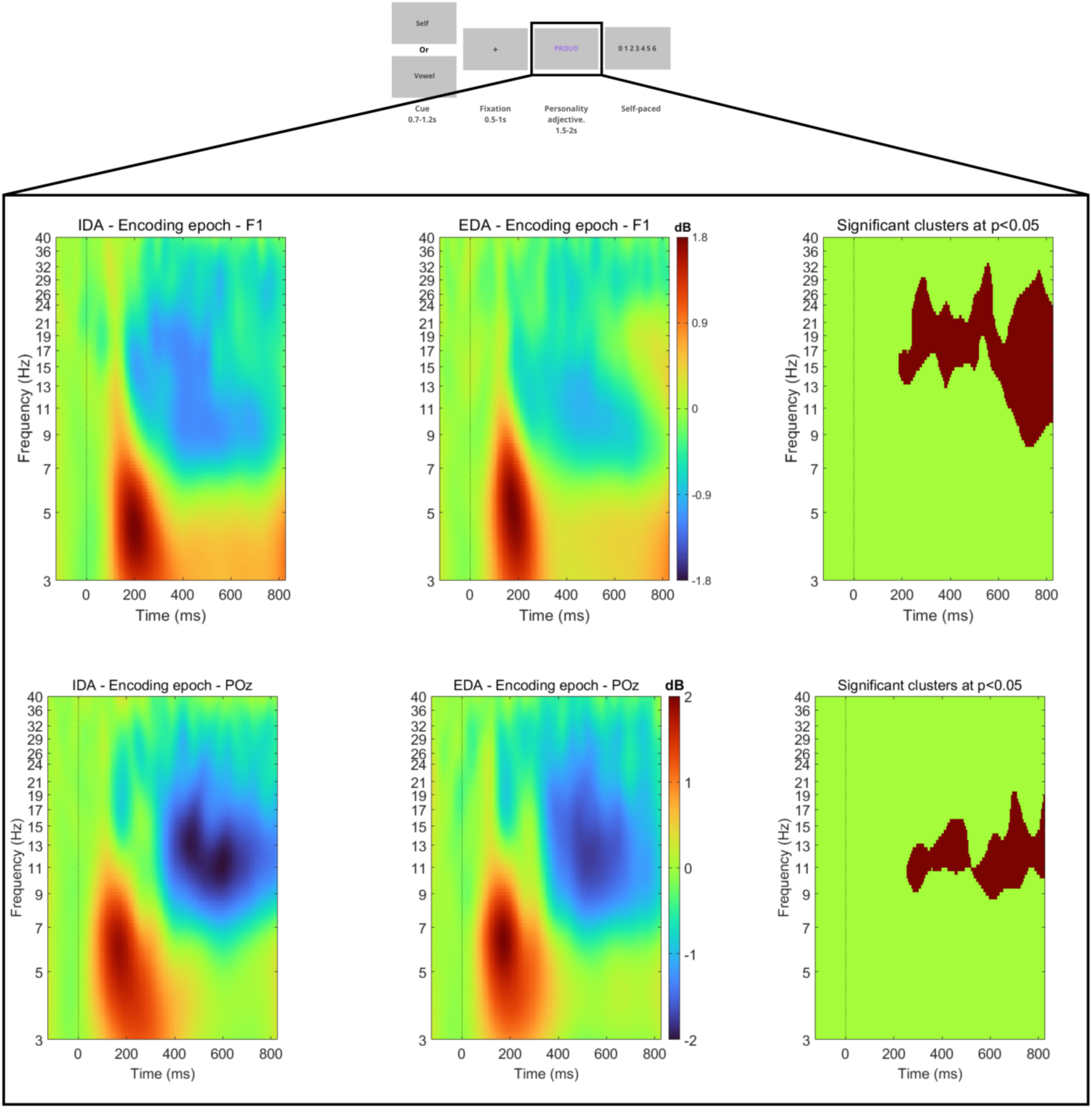
Time-frequency representations during the encoding epoch at frontal (F1; top row) and parietal (POz; bottom row) electrodes separately for each condition. Left and middle panels show power changes (in dB) for the Internally Directed Attention (IDA) and Externally Directed Attention (EDA) conditions, respectively. Right panels show significant time-frequency clusters (p < 0.05, cluster-corrected) comparing IDA and EDA. Red regions indicate statistically significant differences between conditions. Power values are baseline-corrected and averaged over trials and participants.

**Supplementary Table 1.** List of personality adjectives sourced from (Anderson, 1968). Based on an earlier study (Davey et al., 2016), words centered around the median ‘likeableness’ rating reported in the original dataset were selected. The adjectives were matched on valence. In the EDA condition, adjectives were restricted to words containing two to four vowels, based on pilot testing to equalize reaction times across conditions and maintain a consistent overall delay duration (see Supplementary Figure 1).

| **Adjectives in IDA** |  | **Adjectives in EDA** |  |
| --- | --- | --- | --- |
| SILLY | DAYDREAMER | CRITICAL | UNHAPPY |
| SELFISH | CALM | UNHEALTHY | STUBBORN |
| OPPORTUNIST | UNPOPULAR | SARCASTIC | ALERT |
| IDEALISTIC | MEDIOCRE | CONFIDENT | PATIENT |
| EMOTIONAL | SUPERSTITIOUS | RESERVED | MODEST |
| OPTIMISTIC | ADVENTUROUS | RELAXED | INSECURE |
| CRAFTY | WORRYING | UNLUCKY | UNTIDY |
| SELF-CONSCIOUS | BOLD | DARING | LONELY |
| UNTRUTHFUL | PROGRESSIVE | DEPENDENT | OUTGOING |
| ANGRY | AGGREABLE | PROUD | CHOOSY |
| IMPOLITE | IMPULSIVE | ETHICAL | CURIOUS |
| CLEAR-HEADED | SELF-CRITICAL | QUIET | FEARLESS |
| ANNOYING | UNPREDICTABLE | PRACTICAL | QUICK |
| ABSENT-MINDED | PERSUASIVE | HOPEFUL | FORGETFUL |
| TALKATIVE | ENTERTAINING | AVERAGE | MORAL |
| NEAT | UNOBSERVANT | GREEDY | SYSTEMATIC |
| SHY | REALIST | BORING | STUDIOUS |
| PERFECTIONISTIC | CLUMSY | PERSISTENT | CONSISTENT |
| DISHONEST | SELF-CONFIDENT | SILENT | CHILDISH |
| CAUTIOUS | UNCONVENTIONAL | ANXIOUS | EASYGOING |
| UNFRIENDLY | POSSESSIVE | NERVOUS | MOODY |
| AGGRESSIVE | TIDY | TENSE | FEARFULL |
| INCONSISTENT | UNEMOTIONAL | TOUGH | TALENTED |
| NARROW-MINDED | SERIOUS | WORRIER | DIRECT |
| COURAGEOUS | HUMBLE | HESITANT | ABUSIVE |
| AGRUMENTATIVE | OVERSENSITIVE | NOISY | AIMLESS |
| REBELLIOUS | OBEDIENT | ORDERLY | EXCITED |
| UNPUNCTUAL | STRICT | CHARMING | CRUEL |
| UNIMAGINATIVE | LAZY | SKILLFUL | RESTLESS |
| MATERIALISTIC | SENTIMENTAL | UNFAIR | CAREFUL |

## List of Abbreviations

IDA: Internally directed attention
EDA: Externally directed attention
EEG: Electroencephalography
LPP: Late positive potentials
SRP: Self-referential processing
DMN: Default mode network
ERP: Event related potentials
ICA: Independent component analysis
SVT: Simple voltage threshold
MW: Moving window
RT: Reaction time
AIC: Akaike information criterion
ERD: Event related desynchronization
LQMM: Linear quantile mixed models
mPFC: Medial pre-frontal cortex
SD: Standard deviation
SE: Standard error

## Notes

### Competing Interest Statement

The authors have declared no competing interest.

